# Environmental variables determining the distribution of an avian parasite: the case of the *Philornis torquans* complex (Diptera: Muscidae) in South America

**DOI:** 10.1101/839589

**Authors:** Pablo F. Cuervo, Alejandro Percara, Lucas Monje, Pablo M. Beldomenico, Martín A. Quiroga

## Abstract

*Philornis* flies are the major cause of myasis in altricial nestlings of neotropical birds. Its impact ranges from subtle to lethal, being of major concern in endangered bird species with geographically-restricted, fragmented and small-sized populations. In spite of its relevance for bird conservation, there is little information about the environmental dimensions determining their geographical range. We identified for the first time the macro-environmental variables constraining the abiotic niche of the *P. torquans* complex in South America, and provided a model map of its potential distribution based on environmental suitability. We used maximum entropy to model the ecological niche of the *P. torquans* complex, and to determine the contributing scenopoetic variables. We applied most of the recommendations to produce robust ecological niche models: accounting for sampling bias and spatial autocorrelation; using an accessible area for model calibration; testing multiple combinations of model parameters; using multiple statistical criteria; using a number of model replicates to determine consistency and uncertainty in geographical predictions. We identified the minimum temperature of the coldest month as the most relevant variable, associated with the largest decrease in habitat suitability in Brazil and northern South America. Furthermore, the mean temperature of the warmest quarter limited suitability mostly along the Andean range. In addition, humidity and moisture are influential factors in most of Argentina, northern Chile and coastal Peru. The geographical projection of the niche model suggests that environments in most of central-eastern Argentina, and in a broad area in central Chile, are suitable for the presence of the *P. torquans* complex. Besides of contributing to the knowledge of the ecology of the genus, this study is of relevance as a tool for bird conservation and represents a good reference for future work on the distribution of this parasite genus.

## Introduction

In the Neotropics, *Philornis* Meinert 1890 [Diptera: Muscidae] is the major cause of myasis in birds (Guimarães & Papavero, 1999; Dudaniec & Kleindorfer, 2006). *Philornis* adults are free living, while larvae are largely associated with altricial nestling birds, establishing two types of parasitic associations: semi-haematophagous or subcutaneous (Teixeira, 1999). The impact of *Philornis* spp. on hosts ranges from subtle (Dudaniec & Kleindorfer, 2006) to strong and lethal (Dudaniec *et al.*, 2007; Quiroga & Reboreda, 2012), being of particular concern in vulnerable and endangered bird species (Bulgarella *et al.*, 2019). While in healthy populations effects of nest parasitism could sometimes be negligible, in geographically-restricted, fragmented and small-sized populations, the effects of *Philornis* parasitism could be exerting additional pressure (Bulgarella *et al.*, 2019); thus, pushing these populations to the edge of extinction. Evidence that these parasitic flies can be a serious threat to bird conservation is the severe impact caused by *Philornis downsi* [Dodge & Aitkens 1968] on Darwin’s finches (i.e. Kleindorfer & Dudaniec, 2016) and by *P. pici* [Macquart, 1854] on Ridgway’s hawks (*Buteo ridgwayi* - Hayes *et al.*, 2018). Indeed, it may cause up to 75% mortality of nestlings (Fessl *et al.*, 2006; Hayes *et al.*, 2018).

Despite this genus comprises about 50 fly species (Teixeira, 1999) and parasitize more than 250 bird species in the Neotropics (Couri, 1999; Dudaniec & Kleindorfer, 2006; Antoniazzi *et al.*, 2011), there are enormous gaps of information about aspects of its biology and ecology (Dudaniec & Kleindorfer, 2006; Patitucci *et al.*, 2017). One aspect of particular interest and poorly known is the environmental dimensions determining the geographical range of *Philornis* spp. A better understanding of the distribution of this genus may be of relevance to identify locations with high presence of these parasitic flies that might threaten populations of endangered bird species. Furthermore, it may be also appropriate to locate habitats with low risk of *Philornis* occurrence, useful for the re-introduction of bird species, the reinforcement of their populations, or to identify *Philornis-free* areas that are prone to be colonized.

The present study is aimed to identify what macro-environmental variables drive the geographical distribution of *Philornis* species in South America by assessing their relative contribution to the environmental niche. We focus on *P. torquans* and *P. seguyi*, a relatively well-known complex of species within *Philornis*, to start understanding the role of scenopoetic dimensions on the geographical distribution of *Philornis.* As temperature and rainfall are presumed to be the most relevant abiotic parameters affecting *Philornis* spp. life cycle (Dudaniec *et al.*, 2007; Antoniazzi *et al.*, 2011; Manzoli *et al.*, 2013), we hypothesise that low temperatures and reduced precipitation will constrain the occurrence of these species. Ultimately, our contribution to the knowledge on the geographical distribution of these parasitic flies might prove a powerful tool relevant for bird conservation all over the neotropics.

## Materials and Methods

### Study species

*Philornis torquans* [Nielsen, 1913] and *P. seguyi* [Garcia, 1952] are the two subcutaneous *Philornis* species most commonly occurring in southern South America (Couri *et al.*, 2009; Silvestri *et al.*, 2011), where eight *Philornis* species have been described but only half of them are recognized as valid by Couri *et al.* (2009) (namely, *P. torquans*, *P. seguyi*, *P. blanchardi* [Garcia, 1952] and *P. downsi).* Specimens identified morphologically as either *P. torquans* or *P. seguyi* are indistinguishable at the molecular level (Monje *et al.*, 2013). This is the reason why it was suggested that the subcutaneous *Philornis* morphs resembling *P. torquans* or *P. seguyi* in southern South America should be treated as *Philornis torquans* complex (Quiroga *et al.*, 2016). This complex of parasitic flies was selected because: i) its biology and ecoepidemiology have been studied for more than a decade (Antoniazzi *et al.*, 2011; Quiroga & Reboreda, 2012; Manzoli *et al.*, 2013, 2018; Saravia-Pietropaolo *et al.*, 2018); ii) this is the group of flies with the highest number of confirmed morphology-based records; and iii) it is known to affect at least 34 species of birds (see Table S1), and suspected to parasitize endangered species, such as the Saffron-Cowled Blackbird (*Xanthopsar flavus* - BirdLife International, 2019) and the Yellow Cardinal (*Gubernatrix cristata*-BirdLife International, 2018). Both bird species were reported as hosts of unidentified *Philornis* species (Domínguez *et al.*, 2015; Pucheta *et al.*, 2019) in the vicinity of *P. torquans* complex records.

### Species occurrence data

Occurrence data of *Philornis* species was obtained from two different sources: i) field surveys conducted in Argentina; and ii) a thorough review of existing literature. Regarding field surveys, *Philornis* larvae and pupae were recovered from hand-inspected nestlings (during the breeding season) or disassembled nest remains (year-round) and reared to adulthood in captivity by AP. Adults were identified to species level by MAQ using the available taxonomic key (Couri, 1999; Couri *et al.*, 2009). Our literature survey included 265 peer-reviewed publications ranging from 1853 until 2019, which we then systematically reviewed for localities were parasitism occurred. Due to the lack of occurrence data of free-living adults, information obtained from immature *Philornis* stages represents a valid surrogate to assess the environmental suitability for the species.

The specimens identified based on morphology as *P. torquans* or *P. seguyi* were considered as members of the *Philornis torquans* complex (see Quiroga *et al.*, 2016), which accounted for 80 occurrence records (most of them concerning different host species in the same site). After removing duplicate localities, the dataset was reduced to 34 occurrence sites (Table S1). Since spatial autocorrelation in environmental information and consequent non-independence of occurrence data could bias predictions (Boria *et al.*, 2014), we spatially filtered sites preserved for calibration with the package “spThin” in R Statistical Software 3.5.0 (Aiello-Lammens *et al.*, 2015; R Core Team, 2018). Spatial thinning of occurrence records provides an easy-to-implement and relatively straightforward method to alleviate the effects of sampling bias (Boria *et al.*, 2014; Aiello-Lammens *et al.*, 2015). To attain a uniform distribution of occurrences and to avoid excessive clustering within any particular region, we explored different filtering distances in sequential steps, visually inspecting results. Once a satisfactory spatial distribution of occurrence data was obtained (i.e., when spatial clustering of records was no longer evident), we ended up with 18 sites for the *P. torquans* complex (Fig. 1). We split the *P. torquans* complex locations randomly in equal quarts: one set of locations for calibration (75%) and another for evaluation during model calibration (25%).

**Figure 1.**
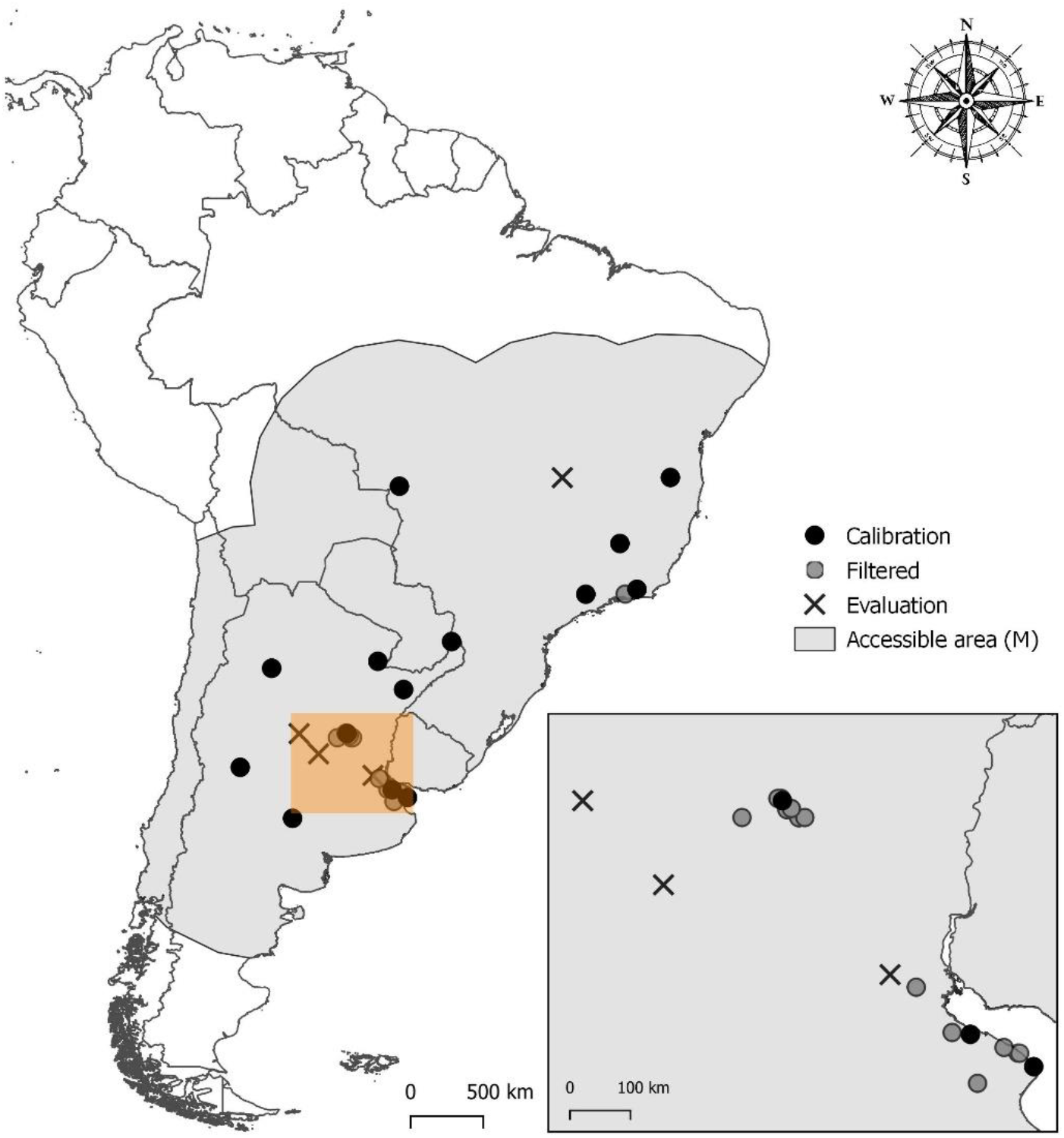
Geographical coordinates representing locations of *Philornis torquans* complex and accessible area (M) used in the construction of models.

### Environmental data and selection of variables

To characterize climatic conditions meaningful to biological species, we used the so-called “bioclimatic” variables based on WorldClim from the CHELSA database (www.chelsa-climate.org, Karger *et al.*, 2017). In addition, we included the climatic variables from the ENVIREM database, derived from and complementing the bioclimatic variables from WorldClim (www.envirem.github.io, Title & Bemmels, 2018). We excluded nine data layers (detailed in supplementary Table S2), as these variables include artefacts that create abrupt differences between neighbouring pixels. We coarsed environmental data to 10 arc minutes (~17 x ~17 km) spatial resolution, which approximately matches the uncertainty inherent in the occurrence data.

As a first step and in order to test our hypothesis, we selected a set of five variables based on current knowledge on the biology of *Philornis* spp.: i) mean temperature of warmest quarter (Bio10); ii) minimum temperature of coldest month (Bio06); iii) precipitation of driest month (Bio14); iv) precipitation of wettest quarter (Bio16) (the four of them from CHELSA); v) CMI, a climatic moisture index from ENVIREM. The “mean temperature of warmest quarter” (Bio10) was selected because this variable is related to the sum of degree days available for development, a usual method used to predict the life stages of other dipteran species (e.g., Psychodidae, Stratiomyidae, Calliphoridae) (Kasap & Alten, 2005; Harnden & Tomberlin, 2016; Yanmanee *et al.*, 2016). The “minimum temperature of the coldest month” (Bio06) was selected due to its negative effect on developmental times and survival of pupae and adults of a wide variety of insects (Sinclair, 2015), therefore arguably limiting *Philornis* distribution. Furthermore, Manzoli *et al.* (2013) found that preceding increases in rainfall were highly correlated with the abundance of *Philornis torquans* parasitic stages. This aspect was considered by including the “precipitation of the driest month” (Bio14) and the “precipitation of the wettest quarter” (Bio16). The “climatic moisture index” (CMI) was included as an integrative marker of environmental conditions, as it is a metric of potential water availability imposed solely by climate (ranges from –1 to +1, with wet climates showing positive values, and dry climates negative (Vörösmarty *et al.*, 2005)). Additionally, we created six other sets of variables to be considered in the evaluation and calibration of models. The variables from the original set of data were grouped in three different sets: i) a dataset including only the 15 bioclimatic variables from CHELSA (referred to hereafter as the *bioclim* dataset); ii) a dataset using the *bioclim* dataset plus the 11 climatic ENVIREM variables (hereafter referred to as the *climatic* dataset); and iii) a dataset using the *climatic* dataset plus the two topographic variables from ENVIREM (hereafter referred to as the *environmental* dataset). Then, we reduced model complexity and collinearity by selecting two simpler sets of variables from each of these three datasets, based on the calibration of preliminary Maxent models (details below). For the selection of these simpler sets of variables, we applied two procedures available in the R package “SDMtune” (Vignali *et al.*, 2019): removal of highly correlated variables (*R* > 0.7), and removal of variables with low importance for the model performance (percent contribution < 5%). Finally, for the calibration of models, we retained six sets of environmental variables: i) a set based on current knowledge on the biology of *Philornis* spp. (Set 1); ii) three sets of uncorrelated variables from the *bioclim, climatic* and *environmental* datasets (Sets 2a, 3a and 4); and iii) two sets of relevant variables selected from the *bioclim* and *climatic* datasets (Sets 2b and 3b) (the selection from the *climatic* and *environmental* datasets resulted in two equivalent sets of relevant variables) (Datasets detailed in Table S2).

### Niche modelling and assessment of variables importance

We used Maxent 3.4.1 for developing models (Phillips *et al.*, 2006). Maxent is a machine-learning method that estimates the species potential geographical distribution by finding the probability distribution of maximum entropy (closest to uniform), subject to the constraint of the expected values of the environmental predictors. Maxent was developed to use presence-only data by contrasting presences against background locations (Phillips *et al.*, 2006; Merow *et al.*, 2013), and has shown to outperform other algorithms, even when applied to small data sets like ours (Hernandez *et al.*, 2006; Wisz *et al.*, 2008; van Proosdij *et al.*, 2016).

To minimize the impact of assumptions about absences from areas that are not accessible to the species (Barve *et al.*, 2011), we restricted model calibration to the area to which the species likely had access via dispersal. Considering there is no available information on the dispersal capacity of the *P. torquans* complex, we defined the adequate accessible area (M) as a maximum area within 1000 km of any of the occurrences (Fig. 1), based on: i) the average geographical distance between a centroid point and all the occurrences (~730 km); ii) the high dispersal capacity (Dudaniec *et al.*, 2008) and assisted dispersion of *Philornis downsi* to the Galapagos Islands (Kleindorfer & Dudaniec, 2016); and iii) the dispersal capacity of other Muscidae species (Sabrosky, 1961; Hogsette & Ruff, 1985).

The complexity of models built with Maxent can be adjusted with the inclusion of additional feature classes (i.e. transformations of the original predictor variables) that allow for increasingly complex models, as well as with a regularization multiplier that contributes to select those features and reduces overfitting (Warren & Seifert, 2011; Merow *et al.*, 2013). The preliminary models for each of the three data sets (the *bioclim, climatic* and *environmental* datasets), later used for variable selection, were calibrated using the “kuenm” R package (Cobos *et al.*, 2019). To determine the optimal model complexity, we explored all combinations of [a] 17 values of the regularization parameter (0.1–1.0 at intervals of 0.1, 2–6 at intervals of 1, and 8 and 10), and [b] nine sets of potential combinations of four feature classes: linear “l”, quadratic “q”, product “p”, and hinge “h” (“l”, “lq”, “lp”, “lqp”, “h”, “lh”, “lqh”, “lph”, “lqph)”.

After the selection of variables, we conducted a detailed model selection exercise, exploring all combinations of [a] the 17 values of the regularization parameter, [b] the nine sets of potential combinations of four feature classes, and [c] the six sets of bioclimatic variables (see above). As such, a total of 918 models were calibrated, and each model was evaluated for statistical significance (partial ROC tests), performance (omission rate), and the Akaike Information Criterion corrected for small sample sizes (AICc). Final models were selected under the three criteria (first, statistical significance, then omission rate, followed by AICc) (Cobos *et al.*, 2019).

We used the best parameter settings chosen to create final models. For each final model, we ran 30 bootstrap replicates, with no clamping or extrapolation, and retaining a random partition of 25% of the points from each run. These final models, though only calibrated across M, were transferred to the extent of South America, particularly to southern South America, which is the focus of this contribution. We used the cloglog output format to assess median values across replicates as an estimate of the spatial distribution of suitable and unsuitable conditions for the *P. torquans* complex across South America. The uncertainty in model predictions was estimated using the range in the suitability values (i.e., maximum minus minimum) across the model replicates. To provide an additional assessment on the reliability of our model transfers, we calculated the mobility-oriented parity (MOP) metric (Owens *et al.*, 2013) to offer a view of the novelty of environmental values and combinations of values across the transfer area, with special attention paid to areas of strict extrapolation (i.e., transfer areas with values outside the range of climates in the calibration area).

We assessed the importance of each variable to determine the environmental suitability for the *P. torquans* complex with median estimates across replicates of percent contribution, permutation importance and Jackknife analysis. In order to determine the most limiting variables, we used the Maxent limiting factor mapping tool described in Elith *et al.* (2010), and implemented via the package “rmaxent” (Baumgartner *et al.*, 2017). The limiting factors are an interesting insight into the drivers of predictions, and provide a powerful basis for the statement of hypotheses concerning physiological tolerance and aptitude (Elith *et al.*, 2010).

## Results

We assessed 918 models for the final model calibration. Seven hundred and twenty-seven of them were statistically significant as compared with a null model of random prediction. Of such significant models, 137 (~19%) met the omission criterion of < 5%. Of the significant, low-omission models, only three met the AICc criterion of ΔAICc < 2. These three models were calibrated with the set of variables selected based on current knowledge (Set 1), and were variants of the regularization multiplier (Table 1). Models based on current knowledge selection of variables outperformed the ones calibrated with automated selected datasets, as the best of models calibrated with Set 1 presented a ΔAICc = 21.8 when compared with the best model calibrated with other set of variables.

**Table 1.**
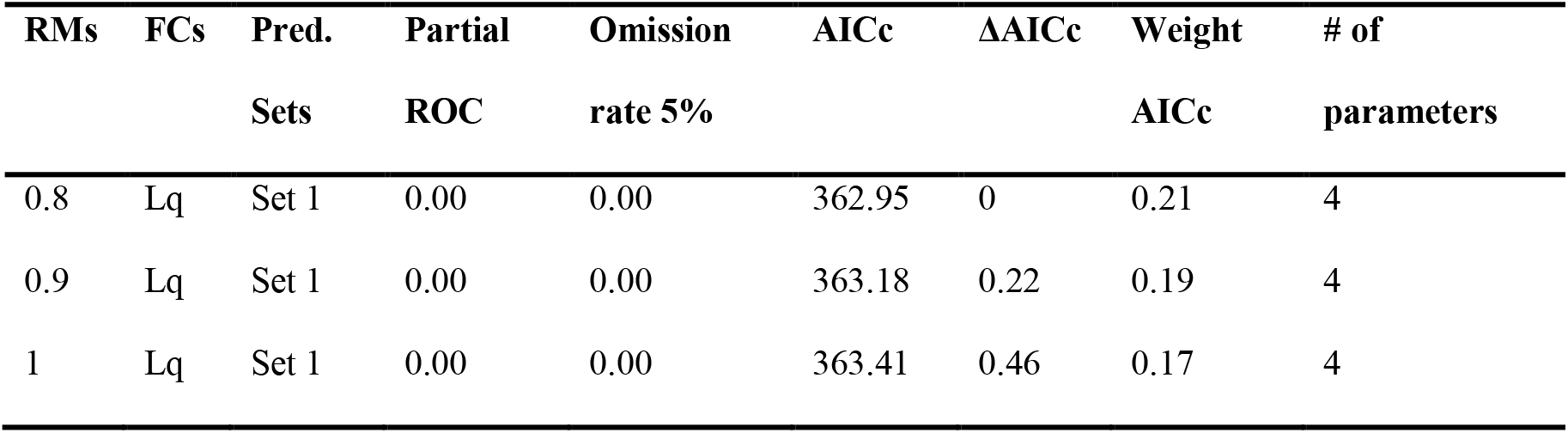
Performance metrics for parameter settings regarding regularization multiplier (RMs), feature classes (FCs), and sets of environmental predictors (Pred. sets), used for creating final models for the *P. torquans* complex. FCs are as follows: linear = l, quadratic = q, product = p, and hinge = h.

The “minimum temperature of the coldest month” (Bio06) was the most influential variable, as evidenced by the percent contribution, permutation importance and Jackknife analysis (Table 2 and Fig. S1). Its response curve had a bell shape, with a central range optimal for prediction, while in both extremes its influence appeared to be detrimental (Fig. S2). The second most important variable was the “mean temperature of warmest quarter” (Bio10), followed by the “climatic moisture index” (CMI). The CMI response curve was similar to the one from Bio06, while the response curve from Bio10 had a sigmoidal shape with positive relation (Fig. S2). The “precipitation of the driest month” (Bio14) and the “precipitation of the wettest quarter” (Bio16) had nearly no influence on the distribution modelling of the *P. torquans* complex.

**Table 2.** Variable importance: percent contribution and permutation importance (median and sd). References: Bio06 = minimum temperature of coldest month; Bio10 = mean temperature of warmest quarter; Bio14 = precipitation of driest month; Bio16 = precipitation of wettest quarter; CMI = climatic moisture index.

| Variable | Percent Contribution | Permutation Importance |
| --- | --- | --- |
| **Bio06** | 54.75 (5.67) | 70.59 (8.09) |
| **Bio10** | 25.10 (5.77) | 18.71 (5.93) |
| **Bio14** | 1.38 (3.29) | 0.64 (4.27) |
| **Bio16** | 0.19 (1.42) | 0.88 (4.43) |
| **CMI** | 12.89 (9.31) | 6.44 (7.07) |

The minimum temperature of coldest month (Bio06) was the predictor associated with the largest decrease in habitat suitability in Brazil and northern South America (Fig. 2). For the majority of Argentina, the climatic moisture index (CMI) was the most limiting factor, while the mean temperature of warmest quarter (Bio10) limited suitability mostly along the Andean range (Fig. 2).

**Figure 2.**
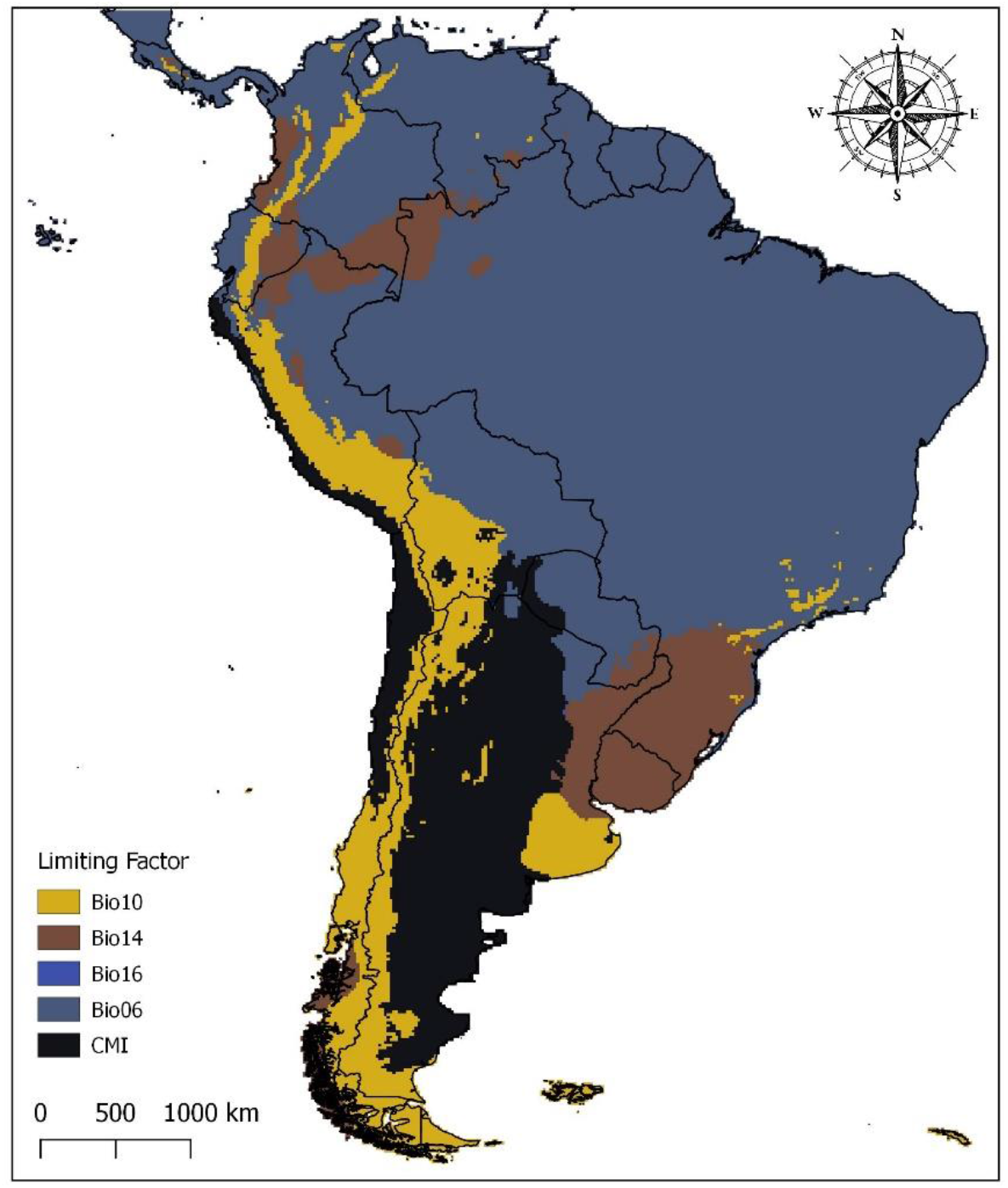
Limiting factors (for any point, the limiting factor is the variable whose value at that point most influences the model prediction). References: Bio06 = minimum temperature of coldest month; Bio10 = mean temperature of warmest quarter; Bio14 = precipitation of driest month; Bio16 = precipitation of wettest quarter; CMI = climatic moisture index. [Colours after the Golden-Collared Tanager *Iridosornis jelskii*]. Patterns of colours for figures were generated with the package “tanagR” (Cadena & Zapata, 2019), inspired by the plumage of passerine birds in the tanager family (Thraupidae) from Central and South America.

As more than one best model was selected, we used the median and range of all replicates across parameters to consolidate results for the *P. torquans* complex. The median of the selected models (Fig. 3A) identified areas with different levels of habitat suitability for the *P. torquans* complex across South America. Highly suitable areas were concentrated on the central-east of Argentina and Uruguay, comprising the Pampean and Chaco (eastern Chaco district) biogeographical provinces (Fig. S3, see in Morrone, 2017). Suitability declined towards western Argentina, southern Brazil and Paraguay. A broad area in central Chile was identified as environmentally suitable, although it may not be accessible to the *P. torquans* complex by natural dispersion. The model showed moderate uncertainty areas mainly throughout central-western Argentina (the Monte biogeographical province), central Chile and the Andean range (Fig. 3B and Fig. S3). The MOP analysis revealed high environmental similarity in most of South America, and thus almost no risk of strict model extrapolation (Fig. 3C).

**Figure 3.**
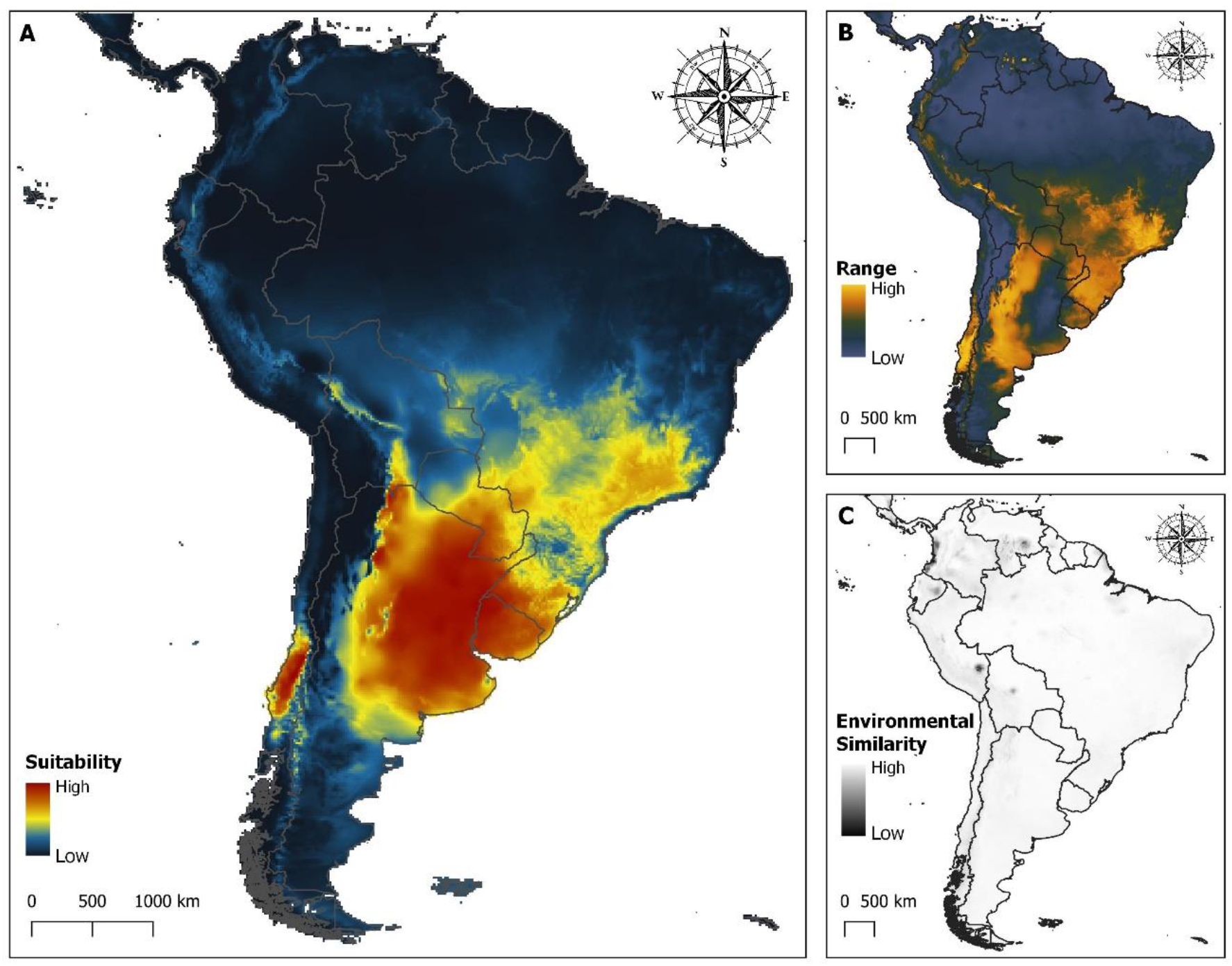
Climatologically suitable regions for the *Philornis torquans* complex distribution in South America. A) Median prediction. B) Uncertainty (range) associated with the median prediction of suitable regions for the *Philornis torquans* complex distribution in South America. C) Mobility-oriented parity analysis revealing similarity of environmental values in South America for the detection of areas of strict extrapolation. [Colours after Scarlet-Breasted Dacnis *Dacnis berlepschi* and Moss-backed Tanager *Bangsia edwardsi*]

## Discussion

In this study, we identified for the first time the macro-environmental (scenopoetic) variables constraining the abiotic niche of the *P. torquans* complex in South America, and provided a model map of its potential distribution based on environmental suitability. Knowledge on the potential distribution of this group of parasitic flies might allow to identify areas where threatened or endemic population of birds may be affected by these parasites; thus being of avian conservation concern.

This study took into consideration most of the recent methodological recommendations to produce robust ecological niche models, as outlined by Fen*g et al.* (2019). As in any modelling approach, model quality is clearly influenced by the number of records used in model building. Thus, it could be argued that the reduced number of occurrences used for modelling in this study is not adequate to produce a reliable niche model. However, Maxent is among the algorithms with best predictive power across a wide range of sample sizes (as low as ten records) (see Hernandez *et al.*, 2006; Wisz *et al.*, 2008), having moderate sample size sensitivity combined with excellent predictive ability (Wisz *et al.*, 2008). Maxent’s strong and consistent performance across sample sizes may be explained by the way it uses regularization to avoid over-fitting (Wisz *et al.*, 2008). The amount of regularization varies flexibly with sample size to ensure consistent performance (Warren & Seifert, 2011; Shcheglovitova & Anderson, 2013), which we have considered when evaluating a number of models across a range of regularization values and features classes (as suggested in Morales *et al.*, 2017). All the same, this modelling approach and its results could benefit from future additions of confirmed occurrence records to the database.

We hypothesised that low temperatures and reduced precipitation will restrain the occurrence of the *Philornis torquans* complex, as these abiotic parameters play a predominant role in establishing the distribution limits of organisms and have already been related with the intensity of parasitism by *Philornis* larvae (Dudaniec *et al.*, 2007; Antoniazzi *et al.*, 2011; Manzoli *et al.*, 2013). Since insects are ectothermic, their rate of development and survival can vary greatly in response to many biotic and abiotic factors (Adams, 1979; Couret *et al.*, 2014). Winters are a critical period for insects at higher latitudes since overwintering conditions strongly affect their fitness (Irwin & Lee, 2000). Indeed, our model identified the minimum temperature of coldest month (Bio06) as the most relevant variable (Table 2 and Fig. S1), supporting our hypothesis. Morphs suggested to belong to the *P. torquans* complex (*P. torquans and P. seguyi*) (Monje *et al.*, 2013) are native to southern South America (with origins somewhere near the border between Brazil and Uruguay - Löwenberg-Neto & De Carvalho, 2009). Representing the southernmost species of the genus, and unlike the majority of *Philornis* species, the *P. torquans* complex must be adapted to endure colder, longer and drier winters, typical of temperate areas.

Furthermore, our model revealed an unexpected detrimental effect in both sides of a suitable range of winter temperatures (signalled by Bio06, Fig. S2), suggesting that the *P. torquans* complex might present a reaction norm of its thermal aptitude along latitudes. Besides the expected negative effect of low temperatures during winter, our model suggests that warm winters may be limiting the northern distribution of the *P. torquans* complex (Fig. 2). Such scenario seems plausible, as insects adapted to cold winters may not perform well in mild winters (Irwin & Lee, 2000), since warmer temperatures may increase energy consumption during this period (Irwin & Lee, 2000; Sinclair, 2015). Therefore, these individuals may not be able to reach the next breeding season or may leave them without enough energy reserves for early reproduction (Sinclair, 2015).

As aforementioned, the developmental rate of insects is related to environmental conditions, particularly as temperature impacts critical rate processes throughout development and adulthood (Kingsolver & Huey, 2008). The temperature dependence of ectotherms’ developmental processes is commonly represented as the cumulative sum of degree-time products required for the development of key life stages, such as egg maturation or emergence from pupae (e.g., Kasap & Alten, 2005; Harnden & Tomberlin, 2016; Yanmanee *et al.*, 2016). In these regards, the mean temperature of the warmest quarter (Bio10) proved to be the second most important variable (Table 2 and Fig. S1), showing a positive relation with sigmoidal shape between predicted probability of presence and Bio10 (Fig. S2). This sigmoidal shape implies a threshold temperature (apparently around 10°C), above which the predicted probability of presence increases exponentially. The former makes sense when considering that the increase of Bio10 might favour the development of temperature-dependent non-parasitic stages, such as pupae maturation, which ultimately relates with the abundance of adults. Otherwise, temperatures below the apparent threshold seems to limit the distribution of the *P. torquans* complex, as evinced along the Andean range (Fig. 2).

The third most relevant variable, the climatic moisture index (CMI), was the most influential factor in most of Argentina, northern Chile and coastal Peru (Fig. 2). The CMI values observed through the actual distribution of the *P. torquans* complex indicate that the species must endure mostly a dry environment when compared with most of South America. As reported for species laying eggs into a relatively dry environment (Grzywacz *et al.*, 2012), the eggs of the *P. torquans* morphs present a small portion of the egg surface that functions as a plastron (Patitucci *et al.*, 2017), a respiratory structure that facilitates oxygen income minimizing water loss (Hinton, 1960). The reduced size of the plastron of the *P. torquans* morph (Patitucci *et al.*, 2017) may have evolved as an adaptation allowing the radiation to the drier conditions typical of central Argentina (see Löwenberg-Neto & De Carvalho, 2009).

Opposed to our prediction, the ecological niche model suggests that precipitation has no direct influence on limiting the environmental suitability of the *P. torquans* complex.

The precipitation of the driest month (Bio14) and the precipitation of the wettest quarter (Bio16) had a poor contribution to the model performance (Table 2), and showed a weak explanatory power in the jackknife analysis (Fig. S1). Though the humidity and moisture are clearly relevant factors determining the *P. torquans* complex distribution, as reflected by CMI and supported by this model, the proximal influence of precipitation seems to be mainly related with the intensity of parasitism by *Philornis* species, as previously reported (Arendt, 1985, 2000; Dudaniec *et al.*, 2007; Antoniazzi *et al.*, 2011; Manzoli *et al.*, 2013).

The outcome of the geographical projection of the environmental niche model was a map of the potential distribution of the *P. torquans* complex in South America. According to our results, the *P. torquans* complex is potentially distributed across central-eastern Argentina, mostly restricted to the Pampean and Chaco biogeographical provinces, and declining with some uncertainty in the surrounding Monte biogeographical province (for more details see in Morrone, 2017) (Fig. 3 and Fig. S3). Despite this niche model is based on scenopoetic variables constrained by accessibility (see Soberón, 2010), but ignoring biotic interactions, the projected potential distribution of the *P. torquans* complex does not seem to be limited by the availability of host species. In fact, the *P. torquans* complex was reported to parasitize at least 34 bird species (Table S1), some of them fairly abundant and widely distributed in South America (e.g., *Pitangus sulphuratus*, *Troglodytes aedon*, *Zonotrichia capensis*). The area predicted as suitable for the *P. torquans* complex in central Chile should be considered with care. First, the estimated suitability is based solely on environmental (scenopoetic) conditions, and thus biotic interactions should prove adequate for the occurrence of the complex in that area. Second, it is required to verify the present occurrence of a *Philornis* species in the predicted area, which could represent either the finding of a new population of the *P. torquans* complex, or the discovery of a sister species. Third, the Andean range might represent a pronounced topographical barrier to be sorted out by the *P. torquans* complex upon its arrival to the estimated suitable area, either by natural dispersion or by human intervention.

In conclusion, this represents the first study to identify macro-environmental variables driving the geographical distribution of a *Philornis* species by evaluating their contribution to its’ environmental niche. Besides of contributing to the knowledge of the ecology of the genus and integrating current methodological recommendations to produce a robust ecological niche prediction, our model is of relevance as a tool for bird conservation and represents a good reference for future work on the distribution of this parasite genus.

## Acknowledgements

We greatly acknowledge the contribution made by Salvador S.A., who kindly provided the geographical coordinates from the sites mentioned in Salvador & Bodrati (2013). We also thank Dra. Bulgarella for the critical reading of this manuscript.

This work was funded by Consejo Nacional de Investigaciones Científicas y Técnicas [PIP11220100100261; PIP 11220130100561CO]; and Universidad Autónoma de Entre Ríos [PIDAC S013805/15]. The funders had no role in study design, data collection and analysis, decision to publish, or preparation of the manuscript.

Version 5 of this preprint has been peer-reviewed and recommended by Peer Community In Ecology (https://doi.org/10.24072/pci.ecology.100049).

## Contribution of authors

PFC and MAQ participated in the design and coordination of the study, and drafted the manuscript; PFC performed the data analysis and modelling; AP carried out field surveys and revised the existing literature; LM and PMB provided financial support and critically reviewed the manuscript. All authors revised the manuscript and gave final approval for publication.

## Conflict of interest disclosure

The authors of this preprint declare that they have no financial conflict of interest with the content of this article

## Supplementary material

Supplementary material freely available in https://doi.org/10.5281/zenodo.3724552

## Data availability

Raw data concerning occurrence localities of the *Philornis torquans* complex and environmental variables is freely available in https://doi.org/10.5281/zenodo.3725944 Detailed explanation and scripts for variable selection available in https://consbiol-unibern.github.io/SDMtune/index.html, and for detailed calibration and construction of ecological niche models available in https://github.com/marlonecobos/kuenm

## Notes

https://doi.org/10.5281/zenodo.3724552

https://doi.org/10.5281/zenodo.3725944

## References

Adams, T.S. (1979) The reproductive physiology of the Screwworm, *Cochliomyia hominivorax* (Diptera: Calliphoridae) II. Effect of constant temperatures on oogenesis. Journal of Medical Entomology, 15, 484–487.

Aiello-Lammens, M.E., Boria, R.A., Radosavljevic, A., Vilela, B. & Anderson, R.P. (2015) spThin: An R package for spatial thinning of species occurrence records for use in ecological niche models. Ecography, 38, 541–545.

Antoniazzi, L.R., Manzoli, D.E., Rohrmann, D., Saravia, M.J., Silvestri, L. & Beldomenico, P.M. (2011) Climate variability affects the impact of parasitic flies on Argentinean forest birds. Journal of Zoology, 283, 126–134.

Arendt, W.J. (1985) Philornis Ectoparasitism of Pearly-eyed Thrashers. I. Impact on Growth and Development of Nestlings. The Auk, 102, 270–280.

Arendt, W.J. (2000) Impact of nest predators, competitors, and ectoparasites on Pearly-eyed Thrashers, with comments on the potential implications for Puerto Rican Parrot recovery. Ornitologia Neotropical, 13–63.

Barve, N., Barve, V., Jiménez-Valverde, A., Lira-Noriega, A., Maher, S.P., Peterson, A.T., et al. (2011) The crucial role of the accessible area in ecological niche modeling and species distribution modeling. Ecological Modelling, 222, 1810–1819.

Baumgartner, J., Wilson, P. & Esperón-Rodríguez, M. (2017) rmaxent: Tools for working with Maxent in R. R package version 0.8.3.9000.

BirdLife International. (2018) Gubernatrix cristata [WWW Document]. The IUCN Red List of Threatened Species. URL http://dx.doi.org/10.2305/IUCN.UK.2018-2. RLTS.T22721578A131888081.en [accessed on 2018].

BirdLife International. (2019) Xanthopsar flavus [WWW Document]. The IUCN Red List of Threatened Species. URL https://dx.doi.org/10.2305/IUCN.UK.2019-3. RLTS.T22724673A153660526.en [accessed on 2019].

Boria, R.A., Olson, L.E., Goodman, S.M. & Anderson, R.P. (2014) Spatial filtering to reduce sampling bias can improve the performance of ecological niche models. Ecological Modelling, 275, 73–77.

Bulgarella, M., Quiroga, M.A. & Heimpel, G.E. (2019) Additive negative effects of Philornis nest parasitism on small and declining Neotropical bird populations. Bird Conservation International, 29, 339–360.

Cadena, D. & Zapata, F. (2019) tanagR: Tanager-Inspired Color Palettes. R package version 0.1.0.

Cobos, M.E., Peterson, A.T., Barve, N. & Osorio-Olvera, L. (2019) kuenm: an R package for detailed development of ecological niche models using Maxent. PeerJ, 7, e6281.

Couret, J., Dotson, E. & Benedict, M.Q. (2014) Temperature, larval diet, and density effects on development rate and survival of *Aedes aegypti* (Diptera: Culicidae). PLOS ONE, 9, e87468.

Couri, M. (1999) Myiasis caused by obligatory parasites. Ia. *Philornis* Meinert (Muscidae). In Myiasis in man and animals in the neotropical region (ed. by Guimarães, J.H. & Papavero, N.). Editora Plêiade, pp. 51–70.

Couri, M.S., Antoniazzi, L.R., Beldomenico, P. & Quiroga, M. (2009) Argentine *Philornis* Meinert species (Diptera: Muscidae) with synonymic notes. Zootaxa, 62, 52–62.

Domínguez, M., Reboreda, J.C. & Mahler, B. (2015) Impact of Shiny Cowbird and botfly parasitism on the reproductive success of the globally endangered Yellow Cardinal *Gubernatrix cristata*. Bird Conservation International, 25, 294–305.

Dudaniec, R.Y., Fessl, B. & Kleindorfer, S. (2007) Interannual and interspecific variation in intensity of the parasitic fly, *Philornis downsi*, in Darwin’s finches. Biological Conservation, 139, 325–332.

Dudaniec, R.Y., Gardner, M.G., Donnellan, S. & Kleindorfer, S. (2008) Genetic variation in the invasive avian parasite, *Philornis downsi* (Diptera, Muscidae) on the Galápagos archipelago. BMC Ecology, 8, 13.

Dudaniec, R.Y. & Kleindorfer, S. (2006) Effects of the parasitic flies of the genus *Philornis* (Diptera: Muscidae) on birds. Emu, 106, 13–20.

Elith, J., Kearney, M. & Phillips, S. (2010) The art of modelling range-shifting species. Methods in Ecology and Evolution, 1, 330–342.

Feng, X., Park, D.S., Walker, C., Peterson, A.T., Merow, C. & Papeş, M. (2019) A checklist for maximizing reproducibility of ecological niche models. Nature Ecology & Evolution, 3, 1382–1395.

Fessl, B., Sinclair, B.J. & Kleindorfer, S. (2006) The life-cycle of *Philornis downsi* (Diptera: Muscidae) parasitizing Darwin’s finches and its impacts on nestling survival. Parasitology, 133, 739–747.

Grzywacz, A., Szpila, K. & Pape, T. (2012) Egg morphology of nine species of Pollenia Robineau-Desvoidy, 1830 (Diptera: Calliphoridae). Microscopy Research and Technique, 75, 955–967.

Guimarães, J.H. & Papavero, N. (1999) Myiasis in man and animals in the Neotropical Region. Editora Plêiade, São Paulo.

Harnden, L.M. & Tomberlin, J.K. (2016) Effects of temperature and diet on black soldier fly, *Hermetia illucens* (L.) (Diptera: Stratiomyidae), development. Forensic Science International, 266, 109–116.

Hayes, C.D., Hayes, T.I., McClure, C.J.W., Quiroga, M., Thorstrom, R.K., Anderson, D.L., et al. (2018) Native parasitic nest fly impacts reproductive success of an island-endemic host. Animal Conservation, 22, 157–164.

Hernandez, P.A., Graham, C.H., Master, L.L., Albert, D.L., Deborah, L. & Albert, D.L. (2006) The effect of sample size and species characteristics on performance of different species distribution modeling methods. Ecography, 29, 773–785.

Hinton, H.E. (1960) The chorionic plastron and its role in the eggs of the Muscinae (Diptera). Journal of Cell Science, s3-101, 313–332.

Hogsette, J.A. & Ruff, J.P. (1985) Stable Fly (Diptera: Muscidae) Migration in Northwest Florida. Environmental Entomology, 14, 170–175.

Irwin, J.T. & Lee, R.E. (2000) Mild winter temperatures reduce survival and potential fecundity of the goldenrod gall fly, *Eurosta solidaginis* (Diptera: Tephritidae). Journal of Insect Physiology, 46, 655–661.

Karger, D.N., Conrad, O., Böhner, J., Kawohl, T., Kreft, H., Soria-Auza, R.W., et al. (2017) Climatologies at high resolution for the earth’s land surface areas. Scientific Data, 4, 170122.

Kasap, O.E. & Alten, B. (2005) Laboratory estimation of degree-day developmental requirements of *Phlebotomus papatasi* (Diptera: Psychodidae). Journal of Vector Ecology, 30, 328–333.

Kingsolver, J.G. & Huey, R.B. (2008) Size, temperature, and fitness: three rules. Evolutionary Ecology Research, 10, 251–268.

Kleindorfer, S. & Dudaniec, R.Y. (2016) Host-parasite ecology, behavior and genetics: A review of the introduced fly parasite *Philornis downsi* and its Darwin’s finch hosts. BMC Zoology.

Löwenberg-Neto, P. & Carvalho, C.J.B. De. (2009) Areas of endemism and spatial diversification of the Muscidae (Insecta: Diptera) in the Andean and Neotropical regions. Journal of Biogeography, 36, 1750–1759.

Manzoli, D.E., Antoniazzi, L.R., Saravia, M.J., Silvestri, L., Rorhmann, D. & Beldomenico, P.M. (2013) Multi-level determinants of parasitic fly infection in forest passerines. PLoS ONE, 8, e67104.

Manzoli, D.E., Saravia-Pietropaolo, M.J., Antoniazzi, L.R., Barengo, E., Arce, S.I., Quiroga, M.A., et al. (2018) Contrasting consequences of different defence strategies in a natural multihost-parasite system. International Journal of Parasitology.

Merow, C., Smith, M.J. & Silander, J.A. (2013) A practical guide to MaxEnt for modeling species’ distributions: What it does, and why inputs and settings matter. Ecography, 36, 1058–1069.

Monje, L.D., Quiroga, M., Manzoli, D.E., Couri, M.S., Silvestri, L., Venzal, J.M., et al. (2013) Sequence analysis of the internal transcribed spacer 2 (ITS2) from *Philornis seguyi* (García, 1952) and *Philornis torquans* (Nielsen, 1913) (Diptera: Muscidae). Systematic Parasitology, 86, 43–51.

Morales, N.S., Fernández, I.C. & Baca-González, V. (2017) MaxEnt’s parameter configuration and small samples: are we paying attention to recommendations? A systematic review. PeerJ, 5, e3093.

Morrone, J.J. (2017) Neotropical biogeography. Regionalization and evolution. CRC Press, Inc.

Owens, H.L., Campbell, L.P., Dornak, L.L., Saupe, E.E., Barve, N., Soberón, J., et al. (2013) Constraints on interpretation of ecological niche models by limited environmental ranges on calibration areas. Ecological Modelling, 263, 10–18.

Patitucci, L.D., Quiroga, M., Couri, M.S. & Saravia-Pietropaolo, M.J. (2017) Oviposition in the bird parasitic fly *Philornis torquans* (Nielsen, 1913) (Diptera: Muscidae) and eggs’ adaptations to dry environments. Zoologischer Anzeiger, 267, 15–20.

Phillips, S.J., Anderson, R.P. & Schapire, R.E. (2006) Maximum entropy modeling of species geographic distributions. Ecological Modelling, 190, 231–259.

van Proosdij, A.S.J., Sosef, M.S.M., Wieringa, J.J. & Raes, N. (2016) Minimum required number of specimen records to develop accurate species distribution models. Ecography, 39, 542–552.

Pucheta, M.F., Patitucci, L.D., Bulgarella, M., Pereda, M.I., Giacomo, A.S. Di & Kopuchian, C. (2019) Primer registro de parasitismo de *Philornis* en pichones de Tordo Amarillo *(Xanthopsar flavus)*. In XVIII Reunión Argentina de Ornitología (RAO). Tandil, Argentina, p. 103.

Quiroga, M.A., Monje, L.D., Arrabal, J.P. & Beldomenico, P.M. (2016) Datos moleculares nuevos sobre *Philornis* (Diptera: Muscidae) subcutáneas del sur de Sudamérica sugieren la existencia de un complejo de especies. Revista Mexicana de Biodiversidad, 87, 1383–1386.

Quiroga, M.A. & Reboreda, J.C. (2012) Lethal and sublethal effects of Botfly *(Philornis seguyi)* parasitism on House Wren nestlings. The Condor, 114, 197–202.

R Core Team. (2018) R: A language and environment for statistical computing.

Sabrosky, C.W. (1961) Our First Decade with the Face Fly, *Musca autumnalis*. Journal of Economic Entomology, 54, 761–763.

Saravia-Pietropaolo, M.J., Arce, S.I., Manzoli, D.E., Quiroga, M. & Beldomenico, P.M. (2018) Aspects of the life cycle of the avian parasite *Philornis torquans* (Diptera: Muscidae) under laboratory rearing conditions. The Canadian Entomologist, 150, 317–325.

Shcheglovitova, M. & Anderson, R.P. (2013) Estimating optimal complexity for ecological niche models: A jackknife approach for species with small sample sizes. Ecological Modelling, 269, 9–17.

Silvestri, L., Antoniazzi, L.R., Couri, M.S., Monje, L.D. & Beldomenico, P.M. (2011) First record of the avian ectoparasite *Philornis downsi* Dodge & Aitken, 1968 (Diptera: Muscidae) in Argentina. Systematic Parasitology, 80, 137–140.

Sinclair, B.J. (2015) Linking energetics and overwintering in temperate insects. Journal of Thermal Biology, 54, 5–11.

Soberón, J.M. (2010) Niche and area of distribution modeling: a population ecology perspective. Ecography, 33, 159–167.

Teixeira, D. (1999) Myiasis caused by obligatory parasites. Ib. General observations on the biology of species of the genus Philornis Meinert, 1890 (Diptera, Muscidae). In Myiasis in man and animals in the Neotropical Region (ed. by Guimarães, J.H. & Papavero, N.). Pleaide/FAPESP, São Paulo, pp. 71–96.

Title, P.O. & Bemmels, J.B. (2018) ENVIREM: an expanded set of bioclimatic and topographic variables increases flexibility and improves performance of ecological niche modeling. Ecography, 41, 291–307.

Vignali, S., Barras, A. & Braunisch, V. (2019) SDMtune: Species Distribution Model Selection.

Vörösmarty, C.J., Douglas, E.M., Green, P.A. & Revenga, C. (2005) Geospatial indicators of emerging water stress: an application to Africa. AMBIO: A Journal of the Human Environment, 34, 230–236.

Warren, D.L. & Seifert, S.N. (2011) Ecological niche modeling in Maxent: the importance of model complexity and the performance of model selection criteria. Ecological Applications, 21, 335–342.

Wisz, M.S., Hijmans, R.J., Li, J., Peterson, A.T., Graham, C.H. & Guisan, A. (2008) Effects of sample size on the performance of species distribution models. Diversity and Distributions, 14, 763–773.

Yanmanee, S., Husemann, M., Benbow, M.E. & Suwannapong, G. (2016) Larval development rates of *Chrysomya rufifacies* Macquart, 1842 (Diptera: Calliphoridae) within its native range in South-East Asia. Forensic Science International, 266, 63–67.

