## Supplementary Material for "Environmental variables determining the distribution of an avian parasite: the case of the *Philornis torquans* complex (Diptera: Muscidae) in South America" (from another version of this preprint: version 3, 2020-03-06)

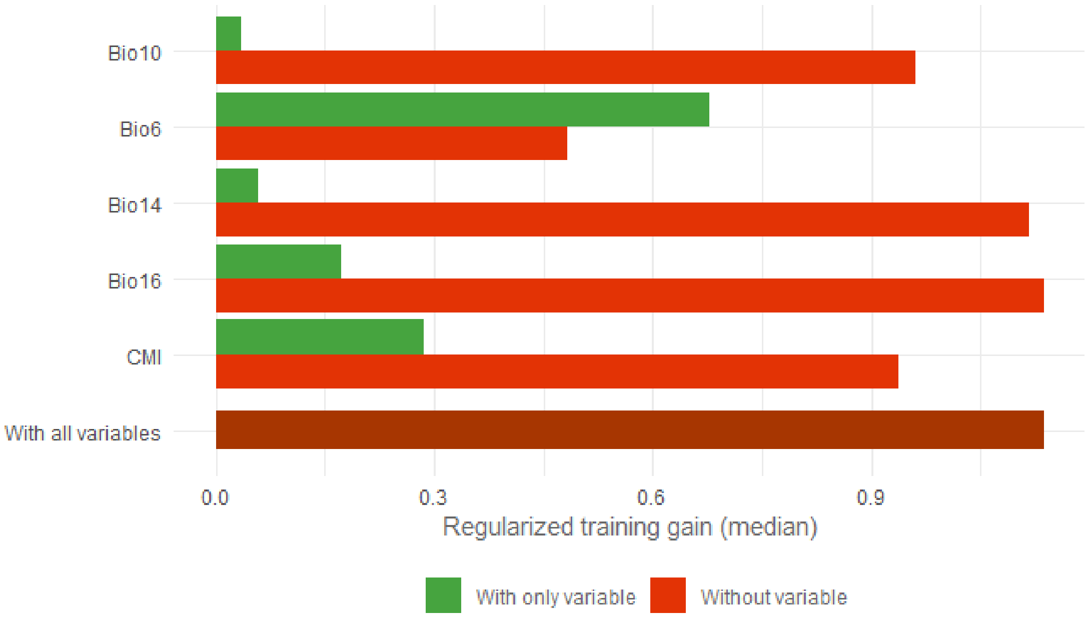


**Figure S1**. Results of the jackknife test of variable importance, when a variable is used in isolation or it is omitted from modelling. References: Bio10 = mean temperature of warmest quarter; Bio06 = minimum temperature of coldest month; Bio14 = precipitation of driest month; Bio16 = precipitation of wettest quarter; CMI = climatic moisture index. [Colours after the Green-Grass Tanager *Chlorornis riefferii*].

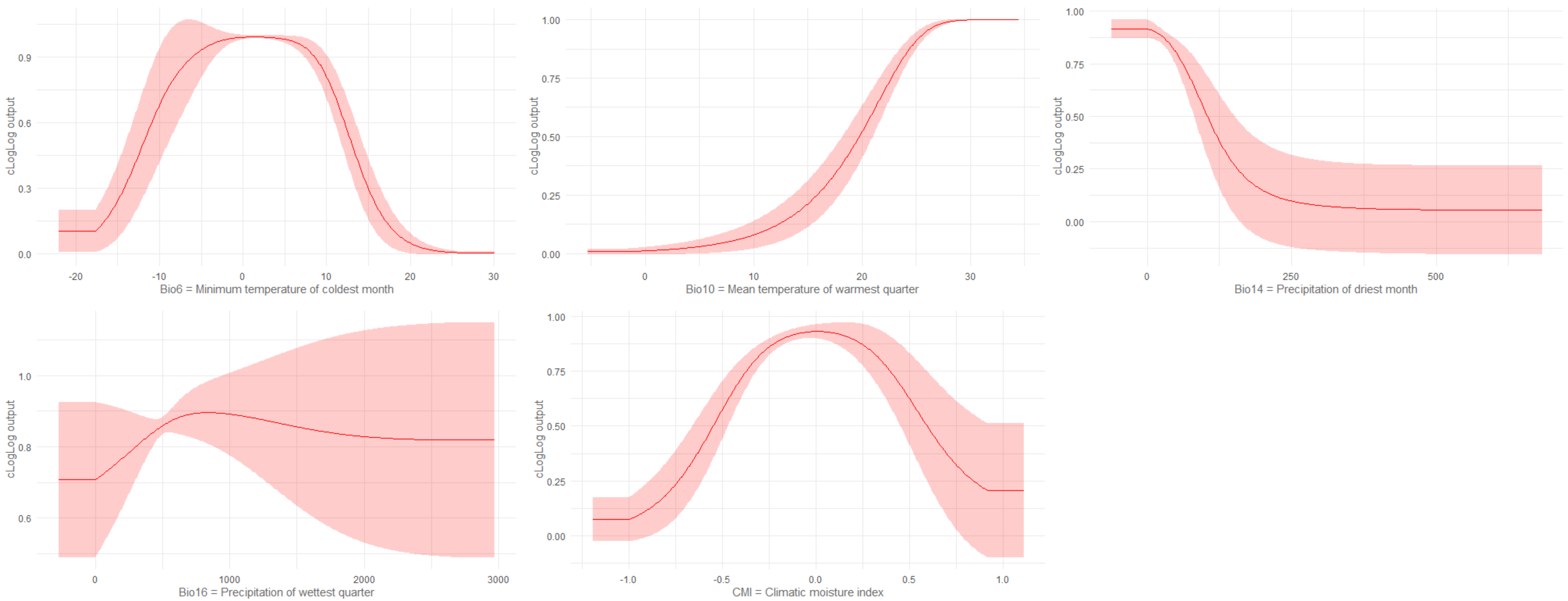


**Figure S2**. Response curves showing how the predicted probability of presence changes as each environmental variable is varied, keeping all other environmental variables at their average sample value.

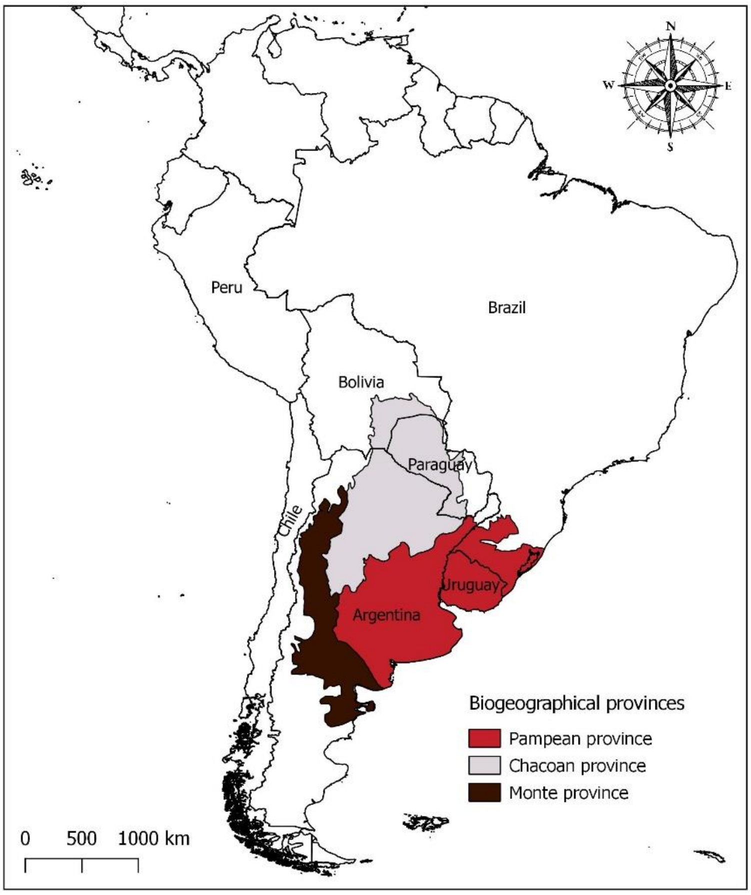


**Figure S3**. Biogeographical provinces from Argentina sustaining environments suitable for the *Philornis torquans* complex (see in Morrone, 2017). [Colours after Multicolored Tanager *Chlorochrysa nitidissima*]

**Table S1**. Geographic coordinates of the presence localities of the *Philornis torquans* complex (morphs *Philornis torquans* and *Philornis seguyi*) included in this study.

| Record N° | Country | Geographic coordinates | Host | References |
|---|---|---|---|---|
| | ***Philornis seguyi*** | | | |
| 1 | Argentina | S 35°34’52”, W 58°1’3” | *Sicalis flaveola* | (Patitucci, 2010) |
| 2 | Argentina | S 35°20’0”, W 57°11’0” | *Agelaioides badius; Mimus saturninus; Paroaria coronata; Troglodytes aedon* | (Segura & Reboreda, 2011) |
| 3 | Argentina | S 35°8’4”, W 57°23’30” | *Anumbius annumbi; Mimus saturninus; Phacellodomus sibilatrix; Phacellodomus striaticollis; Satrapa icterophrys; Sicalis flaveola; Troglodytes aedon; Zonotrichia capensis* | (Mason, 1985) |
| 4 | Argentina | S 35°8’0”, W 57°25’0” | *Mimus saturninus; Molothrus bonariensis; Pitangus sulphuratus; Pyrocephalus rubinus; Satrapa icterophrys; Troglodytes aedon* | (Couri *et al.*, 2005) |
| 5 | Argentina | S 35°2’56”, W 57°37’26” | *Anumbius annumbi; Mimus saturninus* | (Mason, 1985) |
| 6 | Argentina | S 34°51’13”, W 58°7’26” | ND | (Patitucci, 2010) |
| 7 | Argentina | S 34°49’33”, W 58°23’42” | ND | (Patitucci, 2010) |

| | | | | |
|---|---|---|---|---|
| 8 | Argentina | S 34°9’14”, W 58°55’43” | *Anumbius annumbi; Phacellodomus striaticollis; Pitangus sulphuratus* | (Patitucci, 2010) |
| 9 | Argentina | S 33°58’0”, W 59°19’0” | *Mimus saturninus; Molothrus bonariensis* | (Rabuffetti & Reboreda, 2007) |
| 10 | Argentina | S 33°27’56”, W 67°31’55” | *Turdus amaurochalinus* | (Monje *et al.*, 2013) |
| 11 | Argentina | S 31°38’10”, W 60°40’24” | *Troglodytes aedon* | (Quiroga & Reboreda, 2012) |
| 12 | Argentina | S 31°38’0”, W 60°35’0” | *Troglodytes aedon* | (Quiroga & Reboreda, 2012) |
| 13 | Argentina | S 31°38’0”, W 61°31’0” | *Troglodytes aedon* | (Couri *et al.*, 2009) |
| 14 | Argentina | S 31°23’0”, W 63°53’0” | *Anumbius annumbi; Pseudoseisura lophotes* | (Nores, 1995) |
| 15 | Argentina | S 31°21’0”, W 60°59’0” | *Certhiaxis cinnamomeus* | (Couri *et al.*, 2009) |
| 16 | Argentina | S 28°39’17”, W 57°25’48” | *Phacellodomus ruber* | ₴ |
| 17 | Argentina | S 28°39’5”, W 57°25’13” | *Pitangus sulphuratus* | ₴ |
| 18 | Argentina | S 26°54’24”, W 59°0’54” | *Anumbius annumbi* | (Turienzo & Di Iorio, 2014) |
| 19 | Argentina | S 25°41’0”, W 54°27’0” | *Cacicus haemorrhous* | (Fraga, 2011) |
| | ***Philornis torquans*** | | | |
| 20 | Argentina | S 36°37’15”, W 64°17’28” | *Anumbius annumbi* | (Turienzo & Di Iorio, 2014) |
| 21 | Argentina | S 32°38’0”, W 62°41’0” | *Mimus saturninus* | (García, 1952) |
| 22 | Argentina | S 31°31’0”, W 60°51’0” | *Phacellodomus ruber* | (Couri *et al.*, 2009) |
| 23 | Argentina | S 31°30’0”, W 60°47’0” | *Furnarius rufus; Paroaria coronata; Phacellodomus ruber; Phacellodomus* | (Manzoli *et al.*, 2013) |

| | | | | |
|---|---|---|---|---|
| | | | *sibilatrix; Pitangus sulphuratus; Sicalis flaveola* | |
| 24 | Argentina | S 31°23’08”, W 60°55’0” | *Agelaioides badius; Anumbius annumbi; Coryphistera alaudina; Elaenia spectabilis; Furnarius rufus; Leptasthenura platensis; Machetornis rixosus; Mimus saturninus; Molothrus bonariensis; Paroaria coronata; Phacellodomus ruber; Phacellodomus striaticolis; Phacellodomus sibilatrix; Pitangus sulphuratus; Polioptila dumicola; Pseudoseisura lophotes; Saltator coerulescens; Schoeniophylax phryganophilus; Sicalis flaveola; Sublegatus modestus; Suiriri suiriri; Synallaxis frontalis; Taraba major; Troglodytes aedon; Zenaida auriculata* | (De la Peña *et al.*, 2003; Couri *et al.*, 2009; Antoniazzi *et al.*, 2011; Manzoli *et al.*, 2013) |
| 25 | Argentina | S 31°21’0”, W 60°56’0” | *Certhiaxis cinnamomeus* | (De la Peña *et al.*, 2003) |
| 26 | Argentina | S 27°20’45”, W 65°35’34” | *Pitangus sulphuratus; Sporophila nigricollis* | (Nielsen, 1912; Turienzo & Di Iorio, 2007) |
| 27 | Argentina | S 27°19’48”, W 65°34’48” | ND | (Dodge, 1968) |

| | | | | |
|---|---|---|---|---|
| 28 | Brazil | S 22°45’19”, W 46°8’41” | ND | (Löwenberg-Neto & De Carvalho, 2013) |
| 29 | Brazil | S 22°44’38”, W 43°42’27” | ND | (Löwenberg-Neto & De Carvalho, 2013) |
| 30 | Brazil | S 22°27’0”, W 42°58’48” | ND | (Löwenberg-Neto & De Carvalho, 2013) |
| 31 | Brazil | S 19º37’0”, W 44º02’0” | *Phacellodomus rufifrons* | (Costa *et al.*, 2019) |
| 32 | Brazil | S 16°4’14”, W 57°40’44” | ND | (Löwenberg-Neto & De Carvalho, 2013) |
| 33 | Brazil | S 15º32’30”, W 47º36’0” | *Neothraupis fasciata* | (Couri *et al.*, 2018) |
| 34 | Brazil | S 15°31’53”, W 40°54’34” | ND | (Löwenberg-Neto & De Carvalho, 2013) |

ND: No data given regarding host species; ₴: In this paper

**Table S2**. Variables excluded from selection, and sets of variables included in this study.

| Sets of Variables |
|---|
| **Variables excluded from selection** |
| Bio08 = mean temperature of wettest quarter, |
| Bio09 = mean temperature of driest quarter, |
| Bio18 = precipitation of warmest quarter, |
| Bio19 = precipitation of driest quarter, |
| PETColdestQuarter = mean PET of coldest quarter |
| PETWarmestQuarter = mean PET of warmest quarter |
| PETDriestQuarter = mean PET of driest quarter |
| PETWettestQuarter = mean PET of wettest quarter |
| monthCountByTemp10 = number of months with mean temperature > 10°C |
| **SET 1: based on current knowledge** |
| Bio06 = minimum temperature of coldest month |
| Bio10 = mean temperature of warmest quarter |
| Bio14 = precipitation of driest month |
| Bio16 = precipitation of wettest quarter |
| CMI = climatic moisture index |
| **SET 2a: uncorrelated variables from CHELSA** |
| Bio01 = annual mean temperature |
| Bio02 = mean diurnal range |
| Bio04 = temperature seasonality |
| Bio14 = precipitation of driest month |
| Bio15 = precipitation seasonality |
| **SET 3a, uncorrelated variables from CHELSA + ENVIREM** |
| Bio11 = mean temperature of coldest quarter |
| Bio14 = precipitation of driest month |
| Bio15 = precipitation seasonality |
| embergerQ = Emberger's pluviothermic quotient |
| PETseasonality = monthly variability in potential evapotranspiration (PET) |
| **SET 4, uncorrelated variables from CHELSA + ENVIREM (climatic + topographic)** |
| Bio11 = mean temperature of coldest quarter |
| Bio14 = precipitation of driest month |
| Bio15 = precipitation seasonality |
| embergerQ = Emberger's pluviothermic quotient |
| PETseasonality = monthly variability in PET |
| TRI = terrain roughness index |
| **SET 2b, relevant variables from CHELSA** |

Bio11 = mean temperature of coldest quarter
Bio14 = precipitation of driest month

**SET 3b, relevant variables from CHELSA + ENVIREM**
Bio11 = mean temperature of coldest quarter
Bio14 = precipitation of driest month
CONT = continentality

**References**


Antoniazzi, L.R., Manzoli, D.E., Rohrmann, D., Saravia, M.J., Silvestri, L. & Beldomenico, P.M. (2011) Climate variability affects the impact of parasitic flies on Argentinean forest birds. *Journal of Zoology*, **283**, 126–134.

Costa, G.C.A., Koerich, L.B., Pereira, M.H., Dias, L.R., Figueiredo, O., Sant'Anna, M.R. V, Gontijo, N.F., Couri, M.S. & Araujo, R.N. (2019) Occurrence of the parasitic fly *Philornis torquans* on fledglings of the Rufous-throated Thornbird (*Phacellodomus rufifrons*) in Southeast Brazil. *Journal of Wildlife Diseases*, **55**.

Couri, M.S., Antoniazzi, L.R., Beldomenico, P. & Quiroga, M. (2009) Argentine *Philornis* Meinert species (Diptera: Muscidae) with synonymic notes. *Zootaxa*, **62**, 52–62.

Couri, M.S., Barbosa, L., Marini, M.Â., Duca, C. & Pujol-Luz, J.R. (2018) A new host for *Philornis torquans* (Diptera, Muscidae) from the Brazilian Cerrado. *Papéis Avulsos de Zoologia*, **58**, e20185857.

Couri, M.S., Rabuffetti, F.L. & Reboreda, J.C. (2005) New data on *Philornis seguyi* Garcia (1952) (Diptera, Muscidae). *Brazilian Journal of Biology*, **65**, 631–637.

Dodge, H. (1968) Some new and little known species of *Philornis* (Diptera: Muscidae). *Journal of Kansas Entomological Society*, **41**, 155–164.

Fraga, R.M. (2011) Giant Cowbird (*Molothrus oryzivorus*) parasitism of Red-rumped Caciques (*Cacicus haemorrhous*) in the Atlantic forest, northeastern Argentina. *The Wilson Journal of Ornithology*, **123**, 277–282.

García, M. (1952) Las especies argentinas del género *Philornis* Mein., con descripión de especies nuevas (*Dipt. Anthom.*). *Revista de la Sociedad Entomológica Argentina*, **15**, 277–293.

De la Peña, M.R., Beldomenico, P.M. & Antoniazzi, L.R. (2003) Pichones de aves parasitados por larvas de Philornis sp. (Diptera: Muscidae) en un sector de la provincia biogeográfica del Espinal de Santa Fe, Argentina. *Revista FAVE*, **2**, 141–146.

Löwenberg-Neto, P. & De Carvalho, C.J.B. (2013) Muscidae (Insecta: Diptera) of Latin America and the Caribbean: Geographic distribution and check-list by country. *Zootaxa*, **3650**, 1–147.

Manzoli, D.E., Antoniazzi, L.R., Saravia, M.J., Silvestri, L., Rorhmann, D. & Beldomenico, P.M. (2013) Multi-level determinants of parasitic fly infection in forest passerines. *PLoS ONE*, **8**, e67104.

Mason, P. (1985) The nesting biology of some passerines of Buenos Aires, Argentina. *Neotropical Ornithology*, 954–972.

Monje, L.D., Quiroga, M., Manzoli, D.E., Couri, M.S., Silvestri, L., Venzal, J.M., Cuervo, P.F. & Beldomenico, P.M. (2013) Sequence analysis of the internal transcribed spacer 2 (ITS2) from *Philornis seguyi* (García, 1952) and *Philornis torquans* (Nielsen, 1913) (Diptera: Muscidae). *Systematic Parasitology*, **86**, 43–51.

Morrone, J.J. (2017) *Neotropical biogeography. Regionalization and evolution*, CRC Press, Inc.

Nielsen, J.C. (1912) Mydaea anomala Jaenn., a parasite of South-American birds. *Videnskabelige Meddelelser fra Dansk naturhistorisk Forening i Kjøbenhavn*, **63**, 195–208.

Nores, A.I. (1995) Botfly ectoparasitism of the Brown Cacholote and the Firewood-gatherer. *Wilson Bulletin*, **107**, 734–738.

Patitucci, L.D. (2010) Muscidae (Insecta: Diptera) de la provincia de Buenos Aires. Composición específica y estacionalidad.

Quiroga, M.A. & Reboreda, J.C. (2012) Lethal and sublethal effects of Botfly (*Philornis seguyi*) parasitism on House Wren nestlings. *The Condor*, **114**, 197–202.

Rabuffetti, F.L. & Reboreda, J.C. (2007) Early infestation by bot flies (*Philornis seguyi*) decreases chick survival and nesting success in Chalk-Browed Mockingbirds (*Mimus saturninus*). *The Auk*, **124**, 898–906.

Segura, L.N. & Reboreda, J.C. (2011) Botfly parasitism effects on nestling growth and mortality of Red-Crested Cardinals. *The Wilson Journal of Ornithology*, **123**, 107–115.

Turienzo, P. & Di Iorio, O. (2007) Insects found in birds' nests from Argentina. Part I: A bibliographical review, with taxonomical corrections, comments and a hypothetical mechanism of transmission of cimicid bugs. *Zootaxa*, **52**, 1–52.

Turienzo, P. & Di Iorio, O. (2014) Insects found in birds' nests from Argentina. *Pseudoseisura lophotes* Reichenbach, 1853 and *Anumbius annumbi* (Vieillot, 1817) (Aves: Furnariidae), hosts of *Triatoma platensis* Neiva, 1913 (Hemiptera: Reduviidae: Triatominae). *Zootaxa*, **3766**, 1–82.
